# The utility of a closed breeding colony of *Peromyscus leucopus* for dissecting complex traits

**DOI:** 10.1101/2021.08.14.456359

**Authors:** Phillip N Long, Vanessa J Cook, Arundhati Majumder, Alan G Barbour, Anthony D Long

## Abstract

Although *Peromyscus leucopus* (deermouse) is not considered a genetic model system, its genus is well suited for addressing several questions of biologist interest, including the genetic bases of longevity, behavior, physiology, adaptation, and it’s ability to serve as a disease vector. Here we explore a diversity outbred approach for dissecting complex traits in *Peromyscus leucopus*, a non-traditional genetic model system. We take advantage of a closed colony of deer-mice founded from 38 individuals between 1982 and 1985 and subsequently maintained for 35+ years (∼40-60 generations). From 405 low-pass (∼1X) short-read sequenced deermice we accurately imputed genotypes at 17,751,882 SNPs. Conditional on observed genotypes for a subset of 297 individuals, simulations were conducted in which a QTL contributes 5% to a complex trait under three different genetic models. The power of either a haplotype- or marker-based statistical test was estimated to be 15-25% to detect the hidden QTL. Although modest, this power estimate is consistent with that of DO/HS mice and rat experiments for an experiment with ∼300 individuals. This limitation in QTL detection is mostly associated with the stringent significance threshold required to hold the genome-wide false positive rate low, as in all cases we observe considerable linkage signal at the location of simulated QTL, suggesting a larger panel would exhibit greater power. For the subset of cases where a QTL was detected, localization ability appeared very desirable at ∼1-2Mb. We finally carried out a GWAS on a demonstration trait, bleeding time. No tests exceeded the threshold for genome-wide significance, but one of four suggestive regions co-localizes with Von Willebrand factor. Our work suggests that complex traits can be dissected in founders-unknown *P. leucopus* colony mice in much the same manner as founders-known DO/HS mice and rats, with genotypes obtained from low pass sequencing data. Our results further suggest that the DO/HS approach can be powerfully extended to any system in which a founders-unknown closed colony has been maintained for several dozen generations.

## INTRODUCTION

Variation in complex genetic traits is due to the action of many genes as well as the environment. Despite complex genetic traits (*e.g*., risk of certain mental disorders, heart disease, stroke, and diabetes) accounting for the bulk of US health spending, in most cases we do not yet understand their precise genetic architecture. Over the last decade, human geneticists have largely employed a Genome-Wide Association Study (GWAS) approach, using large case-control panels to identify over 70 thousand factors contributing to complex traits (Manolio *et al*. 2009; Buniello *et al*. 2019), with the vast majority of identified factors having extremely subtle phenotypic effects (Boyle *et al*. 2017). In contrast, in major genetic model systems a Multi-Parent Population (MPP) approach has emerged, where quantitative trait loci are mapped in panels derived from multi-generation crosses between several inbred parental lines (de Koning and McIntyre 2017). In the classic MPP-approach to QTL mapping, the MPP population is sampled at the *F_n_* generation in order to derive a large number of Recombinant Inbred Lines (RILs). MPP-RILs are then genotyped at a large number of markers, and investigators need only phenotype the RILs to map a trait. MPP-RILs are available in mice, Drosophila, maize, arabidopsis, *etc.* and have been successfully used to map hundreds of traits (Long *et al*. 2014; de Koning and McIntyre 2017). Despite the success of MPP-RIL approaches, the cost of maintaining a large number of RILs can be high, and RILs are both “lost” and accumulate deleterious mutations over time. In contrast, as the cost of genotyping via short read sequencing continues to plummet we are increasingly seeing the growth of a “Diversity Outbred”/“Heterogeneous Stock” (DO/HS) paradigm where advanced generation MPPs individuals are directly genotyped and phenotyped (Mott *et al*. 2000; Macdonald and Long 2007; Svenson *et al*. 2012). Indeed there are now multiple HS populations maintained in mice (Svenson *et al*. 2012; Woods and Mott 2017) and rats (Hansen and Spuhler 1984) used explicitly for this purpose.

Although mice and rats have excellent DO/HS populations there are many complex traits appropriately studied in other rodent systems. A rodent system of particular significance is “mice” of the *Peromyscus* genus. Although called mice, *Peromyscus spp.* are cricetine rodents, which include hamsters, and are only distantly related to the major murid models of mice and rats. We henceforth refer to *Peromyscus* as “deermice” to avoid this potential confusion. Deermice are the one of the most abundant mammals in North America, and have enjoyed success as a model system for understanding the genetics of adaptation to high altitude, changes in metabolism and immune function during adaptation to urban environments, the genetics of longevity and life history, the genetics of behavioral traits (such as parental care and nest building), and adaptive spatial changes in coat coloration to avoid predation among other traits (beautifully reviewed in (Bedford and Hoekstra 2015)). Different deermice species are also major reservoirs for several tick-borne diseases including Lyme disease, *Borrelia miyamotoi* relapsing fever, the malaria-line protozoan disease babesiosis, and hantavirus (Barbour 2017). The role of *P. leucopus* as the likely primary reservoir for the bacteria that causes Lyme disease (*Borrelia burgdorferi*) and several other tick-borne diseases is analogous to that of bats as reservoirs for SARS coronaviruses and Ebola virus. In fact, *P. leucopus*’s role as the primary reservoir for the bacteria that causes Lyme disease has led to proposals that it be the first mammal considered for natural release gene drive experiments in North America (Najjar *et al*. 2017).

Our previous effort to create infrastructure to strengthen *P. leucopus*’s role as an emerging model system for the study of infectious and other diseases produced a chromosome-length scaffolded 2.45Gb genome assembly, demonstrated how this annotated assembly can empower RNAseq experiments, and identified ∼42 million intermediate frequency SNPs (Long *et al*. 2019). Despite having an annotated UCSC browsable genome, *P. leucopus* does not have a purpose-constructed HS/DO population that can be used for dissecting complex traits like those available in mice and rats. This being said, *P. leucopus* has a closed breeding colony founded from 1982-1985 from 38 wild caught deermice (“Peromyscus leucopus White-footed mouse LL Stock”) that shows great potential for the dissection of complex traits. We speculated that this colony, by virtue of its being closed for ∼60 generations since founding, has many of the properties of the MPPs widely used in model systems. Importantly, we expect the colony to display increased levels of LD relative to natural populations, where useful LD only extends over a few hundred base-pairs (Long *et al*. 2019), and decreased levels of nucleotide variation. These features of the colony should allow missing genotypes to be imputed from low pass sequencing data and genome-wide association scans effectively carried out. Unlike the mouse and rat HS/DO populations, but like other outbred stocks that may exist for other emerging model species, the 38 wild caught deermice “founders” of the *P. leucopus* LL colony are unknown. We show this lack of founder information may minimally impact on our ability to impute SNP genotypes and carry out a GWAS. Instead the main limitation of having “unknown founders” is a reduced ability to identify candidate causative variants (based on the pattern of SNPs private to certain founders) after traits are mapped.

Here we carry out ∼1X per animal low pass sequencing on 405 *P. leucopus* LL colony deermice and use Stitch (Davies *et al*. 2016) to impute SNPs (and haplotypes) for each individual. We validate imputed genotypes using RNAseq data obtained from a subset of genotyped deermice and show imputed SNP genotypes are generally of high quality. We then simulate QTL contributing 5% to variation in a complex trait under three different genetic models and examine the power of both a SNP- and haplotype-based test to detect phenotype genotype associations. Using an appropriate threshold for genome-wide testing, we show that the power to detect a simulated QTL is ∼15-25% largely irrespective of the underlying genetic model simulated and statistical test employed given a study consisting of 300 animals. Despite modest power at this sample size we consistently observe a strong linkage signal at the locations of simulated QTL, suggesting larger sample sizes will greatly increase power, in a manner consistent with that observed in mouse and rat HS/DO studies (c.f. (Gatti *et al*. 2014)). A potenmtially promising observation is that in simulation replicates having a significant hit we can localize the causative gene to ∼2Mb. We finally carry out a GWAS study for a demonstration trait: the time between when a tail is clipped to obtain a small amount of tissue for genotyping and the associated wound appears to cauterize. No regions reached the threshold for significance, but one suggestive region is associated with Von Willebrand Factor.

## MATERIAL AND METHODS

### Animals

Adult female and male outbred *P. leucopus* of the LL stock were obtained from the Peromyscus Genetic Stock Center (PGSC) of the University of South Carolina (PGSC)(Peromyscus Genetic Stock Center, 2017). The LL stock colony was founded with 38 animals captured near Linville, NC between 1982 and 1985 and has been closed since 1985. Sib-sib matings are proscribed, and complete pedigree records are kept. Animals of the LL stock have mitochondria with the same genome sequence (Barbour *et al*. 2019). The gastrointestinal microbiota of the colony animals have been characterized (Milovic *et al*. 2020). Animals were maintained in the AAALAC-accredited U.C. Irvine vivarium with 2-5 animals per cage according to sex and on 12 hours light-12 hours dark lighting schedule, temperature of 21-23° C, humidity of 30-70%, water *ad libitum*, and a diet of 8604 Teklad Rodent (Harlan Laboratories). The study was carried out in accordance with the recommendations in the Guide for the Care and Use of Laboratory Animals of the National Institutes of Health. University of California Irvine protocol AUP-18-020 was approved by the Institutional Animal Care and Use Committee (IACUC).

### Bleeding time assay

The method was a modification of that of Broze et al (Broze *et al*. 2001). The animals were lightly anesthetized with 3% isoflurane by inhalation with 2 L/min flow of oxygen in a small animal veterinary induction chamber. A sterilized, small animal nail clipper (Conair PRO small; item PGRDNCS) equipped with a guard set at 2 mm was used to sever the tail’s tip, and the timer was started. The exposed tissue was briefly touched every 0.5 min with Whatman No. 2 filter paper. The recorded bleeding time was the number of minutes in half minute intervals until further bleeding of the exposed tail tip ceased for at least 0.5 min after the last touch of filter paper. To prevent further bleeding after the animal was returned to its cage Blood Stop Powder (Durvet) was applied. Bleeding times of less than 1.0 minute were not observed. The range of bleeding times was 1.0 to 13.0 min. For 103 animals for which the bleeding time was repeated 2-4 days later the coefficient of determination (*R^2^*) was 0.87. Of the 103, for only 5 (4.9%) was the difference between the two bleeding determinations more than 1.0 min.

### DNA extractions and short read libraries

The 2 mm sample of the excised terminal tail tissue was subjected to extraction with Qiagen’s DNeasy Blood and Tissue Kit. The tissue was first placed in a 2 ml microcentrifuge tube, to which were then added 180 µl of the kit’s Buffer ATL and 30 µl of Proteinase K (Qiagen) at 20 mg/ml. The tube was vortexed and then placed on a shaker at 56° C at 200-220 rpm for up to 12 hr until the skin tissue had dissolved. Following this step the procedure then followed the manufacturer’s instructions. The DNA concentration was determined using a Qubit 2.0 fluorometer with the Qubit dsDNA HS Assay Kit. Short read libraries were prepared in 96-well plates using 1/5^th^ size Illumina Nextera FLEX chemistry reactions, 50ng of gDNA per mouse, and a custom set of 96 unique barcode pairs. We follow the Illumina protocol through the 12-cycle PCR amplification of tagged products, but then post PCR proteinase-K treat each of the 96 samples (to remove polymerase activity), create a 96-plex pool using 2ul of each sample, and then clean-up a 45ul aliquot of the 96-plex reaction following the FLEX protocol. Each 96-plex reaction was run as a PE100 over three HiSEQ4000 lanes, and deplexed using the Illumina software.

### Processing of raw sequencing data

Raw fastq files for 405 animals were aligned to the reference genome using bwa mem (Li 2013) and default settings. We then used STITCH (Davies *et al*. 2016) to impute genotypes for each of the animals with default settings, and `“-- K=8 --nGen=60 --output_haplotype_dosages=TRUE”`. STITCH takes as input the bam files from the alignment step and a list of SNPs at which imputation takes place, here 17.75M biallelic non-repeat-overlapping SNPs, identified from a previous study that carried out sequencing of 36 colony individuals (Long *et al*. 2019). Only SNPs seen in at least two of the 36 colony animals were considered, thus the SNPs of this study are a subset of all SNPs in the colony, with a bias toward more common variants. Furthermore, the quality of SNP imputations would likely have been poorer for singleton SNPs. The `nGen` parameter reflects the number of generations since colony founding (imputation does not seem very sensitive to this parameter). `K` is the number of founder haplotypes in the population. Although the number of founder haplotypes in the colony is 76, which is much greater than the employed 8, increasing `K` greatly beyond 8 seems to result in poorer quality imputations, so we adopted this lower number. STITCH was initially run on all 405 low-pass sequenced individuals (to increase imputation accuracy), although the simulations of this work focus on a subset of 297 animals for which we measured a quantitative trait. STITCH outputs imputed genotypes and haplotype dosages in a vcf file. Genotype and haplotype dosages were extracted per chromosome using the following bcftools queries: `“%POS [%DS{0}]\n”` for genotypes and `“%POS [%HD{0} %HD{1} %HD{2} %HD{3} %HD{4} %HD{5} %HD{6} %HD{7}]\n”` for haplotypes, and wrangled into more tidy/R file with a python script (see software).

### Validation of Stitch genotype calls

Stitch outputs a “dosage” for each imputed SNP, where dosage is a number between 0 and 2 that estimates the number of ALT alleles per individual. In order to validate Stitch dosage calls, we took advantage of six animals for which RNAseq data from spleens was available. RNAseq data from these animals was aligned to the genome using hisat2 (Kim *et al*. 2019) and SNPs were called using GATK (version 4.1.9.0; (McKenna *et al*. 2010). SNPs in the vcf file were filtered using the following vcftools (version 1.10.2; (Danecek *et al*. 2011)) switches `“--min-alleles 2 --max-alleles 2 --min-meanDP 20 --max- missing 1 --maf 0.15 --remove-indels --minQ 30”` which resulted in 110,188 SNP calls. We consider these “high quality SNP calls”, as they have an average coverage of 93X, and as a result we expect the associated calls to be quite accurate in most cases. The RNAseq dosages were merged with imputed STITCH dosages, and the per SNP error was calculated as the absolute difference between the dosages. We examined these errors as a function of per sample gDNA sequence coverage.

### Genetic models

A weakness of all genetic simulations is that we do not know the true genetic architecture of complex traits. As a result we explore three alternative models in an attempt to ensure any statistical test is robust to model assumptions. Under all models a single “gene” contributes 5%, random Gaussian environmental variation contributes 50%, and polygenic background variance contributes 45% to a complex trait. Under the first model a single SNP is causative, under the second 10 SNPs in a 50kb window are causative, and under the third all SNPs in a 50kb window are causative. The polygenic background was simulated by randomly choosing 4,166 SNPs per chromosome, adding their dosage values by individual, scaling the resulting sums to z-scores, and then rescaling so the variance was 45%. We simulated 2*100 different causative “genes”, with each gene centered on the random location of 100 SNPs evenly spaced on either chromosome 23 or 19, irrespective of their minor allele frequency. In the case of the single SNP causative model, the dosage associated with each causative SNP was rescaled to account for 5% of variation in a complex trait. For the ten and all SNP models, we defined a 50kb region centered on each of the 100 SNPs and further considered 10 randomly chosen SNPs or all SNPs in the region. In the case of the 10 SNP model, we added the dosages, converted them to z-scores, and rescaled so that they accounted for 5% of variation. In the case of the all SNP model, we scaled the dosages to have unit variance before adding them, and then rescaled so that they accounted for 5% of variation in the trait. Thus, under the all SNP model, rare SNPs effectively had larger effect sizes, whereas under the 10-SNP model, intermediate frequency alleles had larger effects. We simulate the single SNP causative model for historical reasons (as this model is widely simulated), although hundreds of human GWAS studies suggest that single intermediate frequency SNPs accounting for more than 5% of variation in a complex trait are extremely uncommon.

Although we simulated causative loci on both chromosomes 23 and 19, we only carried out scans using markers on chromosome 23. We focus on chromosome 23 as it is one of the smaller *P. leucopus* chromosomes at 46Mb (Long *et al*. 2019), or 2% of the genome. Given the huge number of SNPs in this species, it can be onerous to carry out hundreds of genomewide scans. In the case of causative loci on chromosome 23 and a chromosome 23 scan, hits are used to estimate the true positive rate of an experiment, whereas for causative loci on chromosome 19 and a chromosome 23 scan, hits are used to estimate the false positive rate (and establish a threshold for statistical significance).

### Kinship matrix

The individuals of this study are from a closed colony. Although the colony employs a breeding design to minimize inbreeding, some mice are more closely related to one another, and the genetic constitution of the colony is slowly changing over time. It is common in such situations to employ a kinship matrix and statistical models to control for impact of cryptic relatedness on test statistics. We derived a kinship matrix at every thousandth locus using the eight STITCH-generated haplotype dosages. In order to calculate a kinship matrix we first had to convert the dosage of the *j^th^* individual at the *l^th^* locus for the *h^th^* haplotype into the three genotypic probabilities. To do this, we define:

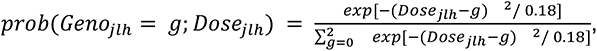

with *g* taking the values 0, 1, or 2 corresponding to the three possible genotypes, and the “0.18” a somewhat arbitrarily chosen constant scaled to reflect uncertainty in those genotypes proportional to the spread of the dosage estimates about 0, 1, or 2. We calculate the proportion of alleles identical by state between individuals *j* and *j’* for the *h^th^* haplotype at the *l^th^* locus as:

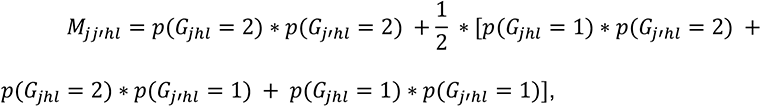

with 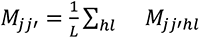 estimating a genome-wide average proportion of alleles IBS. A single kinship matrix was obtained for all 81810 pairs of individuals, and the final matrix (*K*) was multiplied by a constant of 0.79738 so that two parent offspring pairs (not part of the subsetted 297) were forced to have a relatedness of 0.5. We similarly estimated the kinship matrix using every 1000^th^ imputed SNP and the “popkin” package in R (Ochoa and Storey 2021), and observed a correlation coefficient of 0.77 between the two estimates over all pairs of individuals.

### Statistical tests for genotype/phenotype associations

We considered two tests for phenotype and genotype association: a marker- (or SNP-) and haplotype-based test. The marker-based test employs estimated dosages at single SNPs that take on values between 0 and 2 as predictors at any given locus for any given individual. In contrast, the haplotype-based test employs a vector of eight dosages that sum to two, with each element of the vector being the dosage of the particular haplotype (as returned from Stitch). The marker-based scan of chromosome 23 examined 367,487 markers; thus, a complete scan is time-consuming to carry out in the context of a power simulation. In order to carry out all tests without pruning SNPs, we implemented the “fast GWAS” approach of Ziyatdinov (Ziyatdinov *et al*. 2018) that is part of the *lme4qtl* package in R (https://github.com/variani/lme4qtl/blob/master/demo/gwas.R). The scan employs an EVD composition of the “null” model and kinship matrix to enable multiple SNPs to be tested for association in parallel, making the tests very fast. To avoid memory overflows we carried out the scan 5000 loci at a time. Outputed *p*-values were transformed to -log_10_(p-values). Ziyatdinov’s approach approximates fitting the following model with or without the effect of the SNP marker:

*relmatLmer(formula = Y ∼ (1 | ID) + SNP, relmat(ID = K), REML=TRUE)*.

(with *relmatLmer* from the *lme4qtl* package).

We could not employ the same approach for haplotype-based scans and were forced to fit a full model at every locus. We obtained a 10X speed-up by only carrying out a test at every tenth marker. This is a reasonable compromise as haplotype scores change very little over 10 marker intervals, with Supplementary Figure 1 showing that haplotype scores change somewhat slowly over chromosomes. The right side shows this for six individuals using quantiles of haplotype change, and the left side shows haplotype change for two individuals. This figure also shows average haplotype frequencies in the population, typically a small number of haplotypes dominate at any given genomic location. We further sped up testing by creating a new phenotypic variable *Y’* as our dependent variable, obtained by carrying out a principal component analysis on the kinship matrix, regressing the 15 largest principal component scores on *Y*, and retaining the residuals. Finally to avoid problems with co-linear predictors, we also carried out a principal component analysis on the *N*X8 matrix (*H*) of haplotype scores per locus, and retained only principal components scores accounting for greater than 5% of the total variation in the raw haplotype predictor matrix (*H’*). We then fitted the simple linear model to obtain an approximate LOD score:

*anova(lm(Y’ ∼ H’))$’pr(>F)’*

If the LOD10 score for the simple model was greater than 2.0, we fitted the entire mixed model to obtain a more accurate LOD score:

*relmatLmer(formula = Y ∼ (1 | ID) + H’, relmat = list(ID=K), REML=FALSE)*

using *stats::pchisq* and the number of columns in *H’* to compare the mixed model with and without the effect of *H’* to derive a LOD10 score. Fitting every 10^th^ marker, and doing this two step fitting resulted in acceptable but long run times.

### Using false positives to set a significance threshold

For each genetic model and statistical test, we carried out 100 chromosome 23 scans, where a causative QTL was located on either chromosome 23 or chromosome 19. Thus, for any given candidate cut-off threshold, the number of hits associated with a causative locus on chromosome 19 can be used to estimate the false positive rate. Based on quantile-quantile (QQ) plots of causative sites on 19 versus 23 (Figure 2), suitable thresholds do not appear to vary as a function of statistical model, but do vary as a function of statistical test. This is not unsurprising as the marker-based scan employed both a greater number of tests, and tests that in many cases are not as highly correlated with one another. As a result we chose the maximum false positive test statistic over genetic models within statistical tests as a threshold, or 6.0 for the haplotype-based test and 7.5 for the marker-based test. Since chromosome 23 is ∼2% of the genome, and we carried out 100 replicates times three genetic model scans per test, these thresholds hold the genome-wide false single positive rate at approximately one genome-wide false positive per six genome-wide scans.

### Power and localization

A true positive scan is defined as a chromosome 23 scan with a chromosome 23 causative QTL with at least one test statistic greater than the significance threshold. Power is the percent of true positive scans. For each true positive scan we further calculate the total number of markers exceeding the threshold, the distance between the most significant marker (MSM) and the causative SNP (or midpoint of the genetic interval defining a gene), and the distance between the left- and right-most markers above the threshold as the span.

### A demonstration trait

We measured “bleeding time” on 297 of the deermice employed in the above simulations. Each deermouse was also measured for several potential covariates including: <u>Sex</u>, <u>Age</u> at time of assay, <u>Weight</u> at time of assay, and date of birth (<u>DOB</u>; in months from an arbitrary time in the past). Bleeding time is highly skewed with a long tail toward longer times, so the trait was quantile normalized to be normal (qn_Bleed). The model:

*qn_Bleed∼Sex+Age+Weight+DOB*

showed Sex (p<0.021) and Age (p< 0.0007), and DOB (p<1e-10), but not Weight (p<0.2) to be predictive of quantile normalized bleeding time, so we carry out all subsequent analyses on the residuals after Sex, Age, and DOB were removed. Visual examination of plots suggested that Age and DOB affect the trait as first order polynomials. Although bleeding time was measured in colony animals over 6 years, 75% of the animals were assayed over less than two years, but we do not know if our assay was changing slowly over time or the colony itself was changing.

We estimate the heritability, using *lme4qtl::relmatLmer* (below), of the residual quantile normalized bleeding phenotype or a single replicate of our simulation using either the haplotype- or SNP-based kinship matrices described above. In both cases the heritability of the bleeding time phenotype was not different from zero. Positive control heritability estimates employing a single simulated phenotypic replicate (whose expectation is 50%; 45% background plus 5% QTL) resulted in inaccurate heritability estimates, with both kinship matrices resulting in over-estimates of heritability by more than 15%. Since the background genetic variance is constant over relicate simulations we did not attempt to estimate heritability using additional replicate simulations. Although it is not surprising that heritability estimates are inaccurate (given N=300), our estimates do suggest the heritability of bleeding time may be quite low.

We carried out genome wide marker- and haplotype-based scans on quantile normalized bleeding time after Sex, Weight, and DOB were removed, using the significance threshold obtained from the simulations, created Manhattan plots and extracted the top four hits as evidenced from haplotype- and marker-based scores. We furthermore shuffled the residual quantile normalized bleeding time phenotype with respect to the marker data and carried out a second scan, created Manhattan plots, and created QQ-plots of the shuffled LOD scores versus actual LOD scores.

### Platelet function candidate genes

We did an Ensembl Biomart query to extract the house mouse protein sequences associated with the two GO terms associated with platelet function (GO:0030168 and GO:0070527), retained the longest amino acid sequence per gene, blatted those proteins to the *P. leucopus* reference genome, and took the highest scoring hit per gene to obtain the locations of the 83 *P. leucopus* orthologous genes. We consider these platelet function genes to be candidate genes for a GWAS hit if they are within 2Mb of the most-significant marker defining the top four hits. We additionally considered VKORC1 (@chr1:76732685-76735060) and its paralog VKORC1L1 (@chr23:21130849-21176651) as candidate genes. VKORC1 is associated with warfarin resistance in wild rodent populations. Given the 85 candidate genes, the loose 2Mb criteria for a match, and 4 hits tested, we expect ∼1 such hit by chance alone. Thus the observation of a candidate gene within 2Mb of a hit in and of itself is thus **not** strong evidence that that gene is causative.

### Data and software availability

Raw sequencing reads have been uploaded to the SRA under project PRJNA751054 and accessions SAMN20504476 through SAMN20504850 plus a blank (SAMN20504851). We host some useful processed files (SNP and haplotype calls, and individual IDs included in the study) at: http://wfitch.bio.uci.edu/~tdlong/sandvox/publications.html. There is a github archive with code to reproduce analyses: https://github.com/tdlong/Peromyscus_power_bleed.

## RESULTS

### Diploid genotypes can be accurately imputed in colony animals from low pass sequencing data

We carried out low coverage sequencing of 405 colony mice and aligned reads to the *P. leucopus* reference genome (Long *et al*. 2019). Figure 1B, a histogram of observed sequence coverage in the 297 individuals we study in depth in this work, shows that ∼95% of the individuals have coverages between ∼0.5X-2.5X; clearly such coverages are insufficient for directly calling diploid genotypes. We used the program Stitch (Davies *et al*. 2016) to impute genotypes (and eight pseudo-haplotypes) at 17,751,882 previously identified SNPs. In order to validate the genotype calls from STITCH we took advantage of six deer mice for which we additionally obtained RNAseq data from the spleen. We aligned the RNAseq data to the genome, called genotypes, identified a set of 110,188 highly confident genotypes at exonic SNPs, and estimated the absolute error in dosage estimates between imputed genotypes from low coverage data and the highly confident RNAseq calls. Figure 1A is a plot of the percent of the imputed SNPs exhibiting an absolute error greater than 0.10 as a function of the genomic DNA sequencing coverage available to STITCH. It is apparent that at coverages greater than ∼0.3X (*i.e.*, log10(coverage) > -0.5) the quality of imputation is quite high. There is some evidence that the imputation quality improves as coverage approaches 1X, and worsens at much lower coverages (*e.g.,* 0.035X). Figure 1C is a plot of the cumulative proportion of SNPs as a function of their average error rate. We estimate that for a deermouse (in green) sequenced to ∼0.3X, 90% of the imputed SNPs have essentially zero error. But, the majority of individuals have coverages around 1X, where 95% of SNPs have errors of essentially zero. Given the relatively small panel of 405 individuals used in the imputation, and the deermice being part of a closed colony, the quality of these imputations are consistent with claims in the Stitch publication (Davies *et al*. 2016). It is perhaps counter-intuitive that high quality diploid genotypes can be obtained routinely from animals sequenced to <1X, but imputation from low-pass sequencing data appears to be routine in *P. leucopus* closed colony deermice, as there is likely extensive linkage disequilibrium.

**Figure 1:**
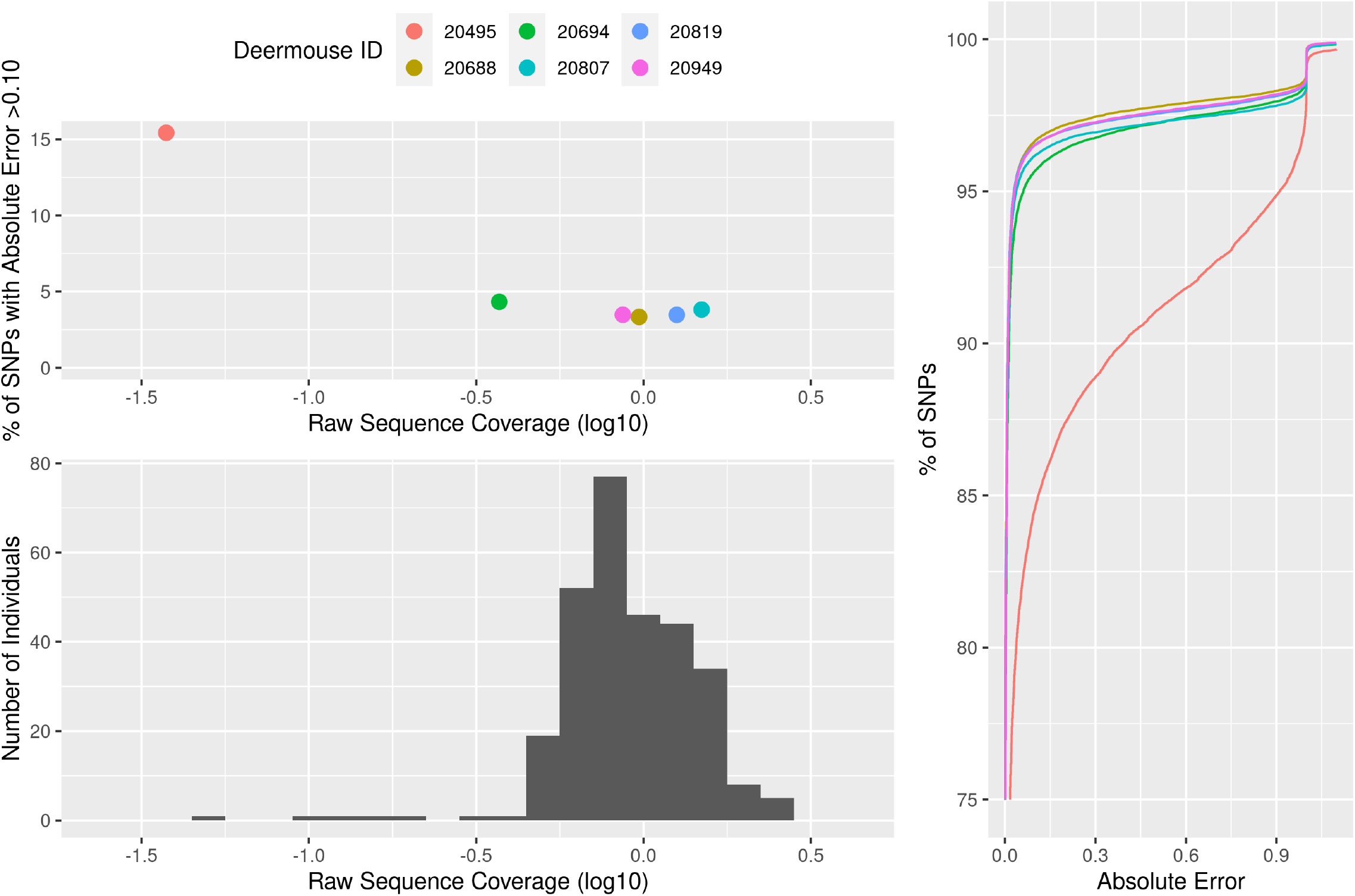
Validation of genotype calls based on six individuals with high coverage RNAseq data. Upper left, percent of SNPs with an absolute error > 0.10 as a function of log10 coverage. Lower left, histogram of raw log10 sequence coverage for the subset of 297 individuals (5 outliers with coverages < 0.025X are not depicted). Right, the cumulative absolute errors.

We estimate a kinship matrix at every 1000th marker genome-wide for the set of 8 haplotype dosages for the 405 animals, and normalized the relatedness matrix using a parent offspring trio (Supp. Figure 2). Although there is the potential for relatedness in the *P. leucopus* colony animals of our study, the kinship matrix subsetted for the 297 individuals we examine more carefully here, suggested that mating between closely related animals is largely avoided in the colony with only a small number of comparisons between individuals suggesting close relatedness. We did not attempt to remove these individuals from the study, and instead utilize the kinship matrix and mixed models to map QTL.

**Figure 2:**
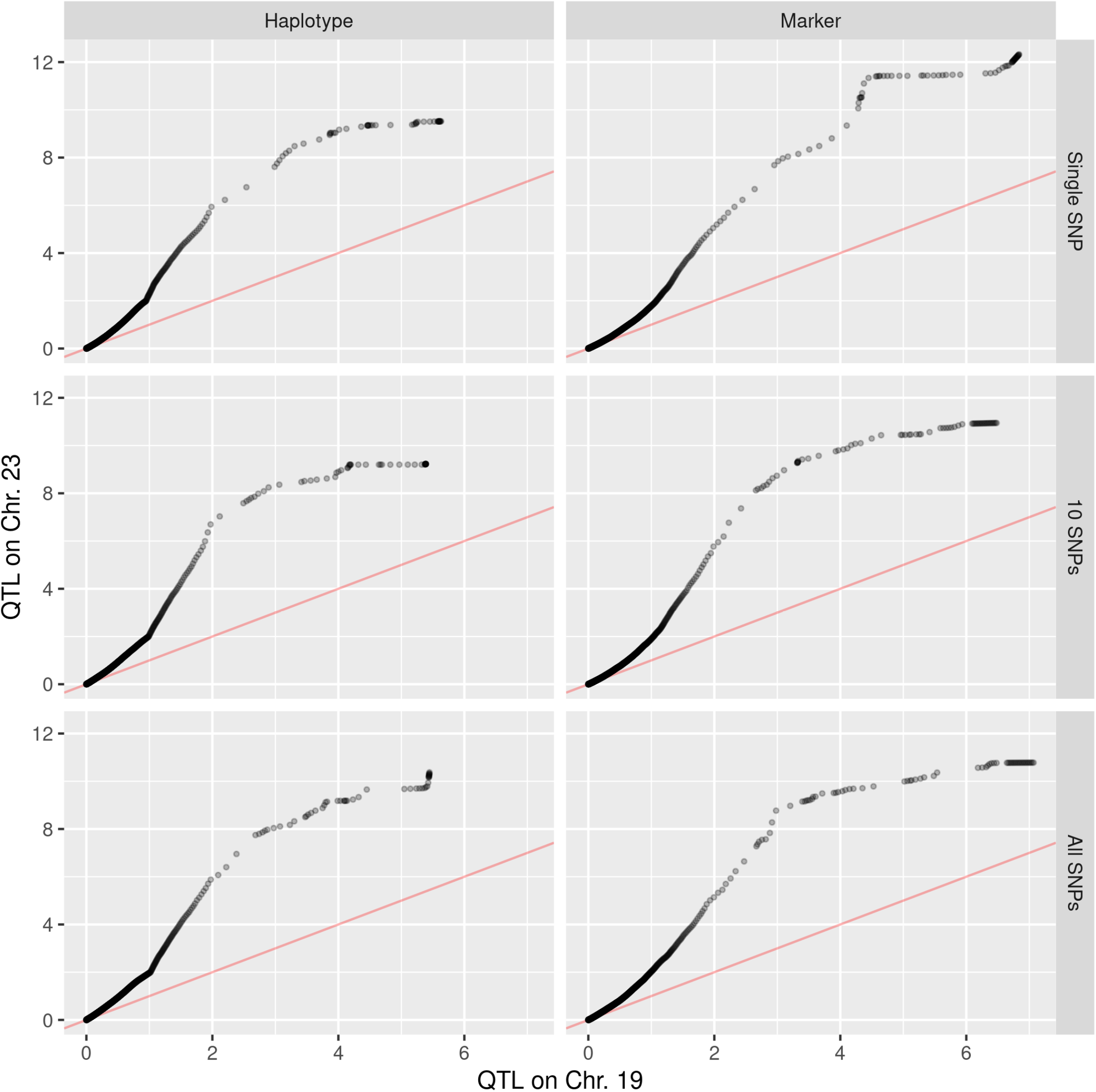
QQ plots of LOD Scores using markers on chromosome 23, with a simulated QTL on either chromosome 23 or 19 (control). Left, haplotype-based tests; right, marker-based tests; top to bottom considers three different genetic models: a single causative SNP, 10 causative SNPs in a 50 Kb “gene” region, or all SNPs in a 50 Kb “gene” region causative. A line with a slope of one is plotted in red.

### Simulations of the power to detect genes contributing to a complex trait in *P. leucopus* colony animals

In order to examine the power of different strategies to detect genes contributing to a complex trait in colony animals, we carried out scans using the 379,282 chromosome 23 SNP markers (or ∼2% of the genome (Long *et al*. 2019)) and a causative gene on either chromosome 23 or chromosome 19. Hits from a chromosome 23 scan, when the causative gene is on another chromosome, allow us to estimate the false positive rate of different tests. Whereas hits from a chromosome 23 scan, where the causative gene is located on the same chromosome, allow us to estimate the true positive rate under a threshold set using the false positive scan. Focusing on a single chromosome made it computationally efficient to examine three different genetic models and two different statistical tests, each with one hundred replicate scans. Even focusing on a single chromosome and employing some simplifications of the statistical models each chromosome 23 scan took 0.5 or 20 CPU hours to run for the marker- and haplotype-based scans respectively, suggesting that entire genome scans would have been computationally prohibitive without other sacrifices. We considered three different genetic models per scan, which briefly included one with a single causative SNP, another with 10 causative SNPs in a 50kb “gene” with equal effect sizes irrespective of minor allele frequency, and a third where all SNPs in a 50kb “gene” are causative and each contributed the same variance to a complex trait. For all models, the causative gene contributed 5% to a complex trait and polygenic background variation 45% to a complex trait with a heritability of 50%. We tested all imputed SNPs for an association with the complex trait using a mixed model approach that leveraged the kinship matrix to control for population structure-based false positives, or a haplotype-based test that employed a set of 8 haplotypes considered internally by Stitch. Like the SNP-based test, the haplotype-based test attempted to control for relatedness; unlike the SNP-based test, the haplotype-based test only considered every tenth position on chromosome 23.

Figure 2 presents QQ-plots by statistical test and genetic model of a chromosome 23 scan for a QTL located on chromosome 23 versus chromosome 19 (the null model) with points representing the product of all tests and 100 replicate simulations. We use the test statistics for the QTL located on chromosome 19 to establish a threshold for statistical significance, 6.0 in the case of the haplotype-based test and 7.5 in the case of the marker-based test. The observation that the threshold required to control for false positives differ between the marker and haplotype-based tests is not unexpected, as the marker-based test employs both more predictors and predictors that are often less correlated with one another than the haplotype-based test. In contrast the thresholds appear to be similar irrespective of the underlying genetic model for the complex trait. Each threshold is established from 100 scans times three genetic models and chromosome 23 is 2% of the genome implying our thresholds result in a false positive rate of less than one false positive per six genome scans. These QQ-plots suggest that there is considerable “signal” when comparing the complex trait gene on chromosome 23 to one located chromosome 19, suggesting that GWAS can work in low pass sequenced *P. leucopus* colony animals given a large enough sample size.

Figure 3 characterizes six realizations of the simulation detecting the same causative “gene” on chromosome 23 (and Supplementary Figure 3 is a similar plot for a control scan with the causative gene on chromosome 19). Although no tests reach the significance threshold level in the control scan there are peaks that could be interpreted as suggestive and tests show a clear block-like pattern (consistent with patterns of LD in the colony). In the example where the causative gene is on chromosome 23, we observe a hit for the 10 causative SNP model under both statistical tests. It is important to note that although the six examples depicted consider the same causative “gene”, the way the genetic models are simulated means that the actual causative nucleotide(s) vary across models, despite all being located in the same 50kb region of the genome (and a second example could be significant under the single SNP or all SNP model). Comparisons between tests within models are valid as the underlying genetics are identical, but comparisons between models are only valid on average. In this example the QTL is well detected and localized by both the haplotype- and marker-based tests, and an experimentalist would be presented with a somewhat defined region to examine for candidate genes that could be causative. The regions displaying the hit appear to display “block-like” significance, and this is typical of many examples. It is noteworthy, and perhaps counter-intuitive, that the marker-based test seems to be similarly powered to detect QTL whose underlying genetics is NOT a single causative site (Table 1). We suspect both tests are really detecting haplotypes or markers that happen to tag a region fairly efficiently (and the marker-based test is not necessarily tagging the causative sites themselves). This is almost certainly true in the cases where the marker-based test has a hit under the all-SNPs model where the effects at individual SNPs are vanishingly small. Finally, although the single-SNP and all-SNP models are not significant in this example they clearly show suggestive peaks. These peaks and the QQ plots of Figure 2 suggest that in an experiment or simulation employing a greater number of individuals these peaks would be much more likely to exceed the threshold and be detected.

**Figure 3:**
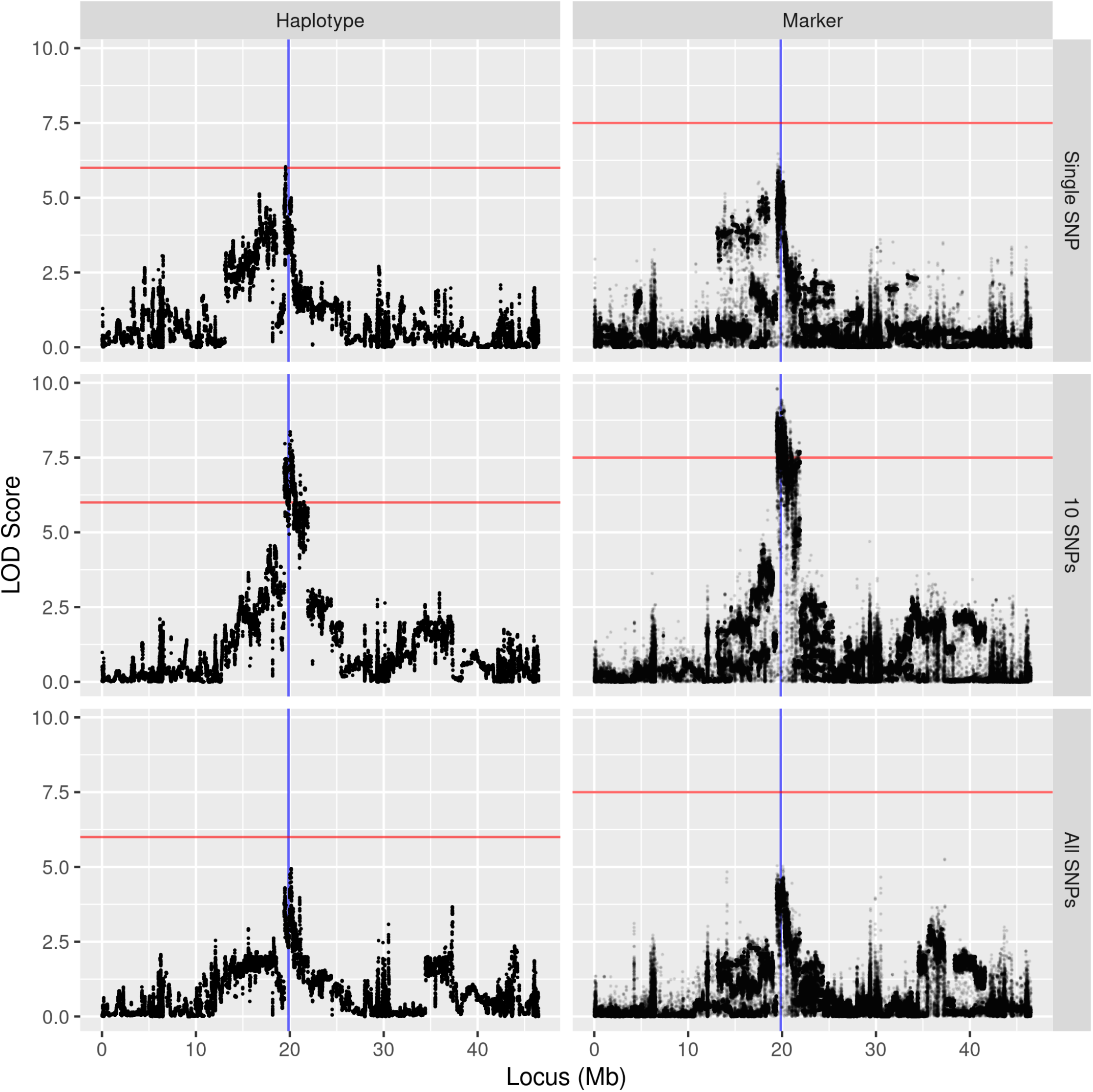
Example of chromosome 23 scans with a simulated QTL on chromosome 23. Left, haplotype-based tests; right, marker-based tests; top to bottom are three different genetic models: single SNP causative, 10 SNPs causative, or all SNPs causative. A red line indicates the significance threshold; a blue line is the location of the causative gene.

**Table 1:** Properties of Chromosome 23 hits

| Genetic Model | Statistical Test | Power (%) * | Given at Least One Hit |  |  |  |
| --- | --- | --- | --- | --- | --- | --- |
|  |  |  | Average # Hits | Average Span (Mb) | Average Distance to MSM (Mb) | Median Distance to MSM (Mb) |
| Single SNP | Haplotype | 29 | 491 | 1.85 | 1.83 | 1.07 |
| Single SNP | Marker | 19 | 2377 | 2.59 | 0.93 | 0.82 |
| 10 SNPs | Haplotype | 25 | 752 | 2.06 | 0.41 | 0.27 |
| 10 SNPs | Marker | 26 | 2690 | 1.92 | 0.54 | 0.38 |
| All SNPs | Haplotype | 31 | 527 | 1.82 | 1.23 | 0.56 |
| All SNPs | Marker | 21 | 1344 | 2.14 | 0.47 | 0.40 |
| Average | Haplotype | 28 | 581 | 1.90 | 1.19 | 0.56 |
| Average | Marker | 22 | 2172 | 2.18 | 0.63 | 0.42 |

\* percent of replicates with at least one test above 6.0 (haplotype) or 7.5 (marker)

We tabulated summary statistics for the scans carried out to detect a QTL located on chromosome 23 (Table 1; thresholds are 7.5 and 6.0 for marker and haplotype-based tests respectively). Interestingly, within the statistical test employed the likelihood of observing at least one hit exceeding our threshold (or power) is largely the same irrespective of the underlying genetic model, with the marker-based approach affording slightly greater power than the haplotype-based approach. This being said, at ∼20% the absolute power is modest. The observation of modest power with N=300 individuals is not surprising, and is consistent with mice/rat DO/HS simulations and studies (Rat Genome Sequencing and Mapping Consortium *et al*. 2013; Gatti *et al*. 2014). It is apparent that dissecting complex traits in *P. leucopus* colony individuals will require a greater number of genotyped and phenotyped individuals. Our threshold was also chosen such that the possibility of a single false positive genome-wide was roughly one in six; employing a more liberal “suggestive” threshold (c.f. Drosophila GWAS (Mackay and Huang 2018)), would increase power at the expense of false positives. We explore four such suggestive peaks below.

For the simulation replicates with at least one marker exceeding the threshold we attempted to quantify localization ability. The average distance between the Most Significant Marker (MSM) and the causative site (or gene mid-point) is consistently less in marker-based tests (0.6 Mb average) than haplotype-based tests (1.4 Mb average). However, the average size of the region above the significance threshold is greater in marker-based tests than haplotype-based tests. Supplementary Figure 4 plots the distance of the MSM from the causative SNP/gene as a function of the LOD score at the MSM. The significance of the MSM is not predictive of localization ability (3rd order polynomial *p*=0.58 and *p*=0.63 for marker- and haplotype-based tests respectively). This lack of a relationship reflects the details of our simulation in part, and if the variance due to the simulated QTL had been allowed to vary, perhaps localization ability would be a function of observed LOD score. On the other hand, it may be that resolution is somewhat constrained by patterns of LD in the colony, as chromosome specific Manhattan plots (c.f. Figure 3) suggest block-like patterns of highly significant markers.

Finally, the median distance of the MSM from the causative SNP/gene is less than the mean distance (Table 1), suggesting that occasionally the MSM is very far from the causative site, Figure 4 examines four such examples. In three of four cases there is considerable signal near the causative QTL, but the largest signal just happened to be elsewhere on the chromosome. This artifact is unlikely to be a pure false positive (as it is not seen when the causative factor is on chromosome 19). Instead we think it could be due to long range LD within a chromosome in the colony animals. We speculate (without proof) that in a larger study the markers near the QTL may eventually exceed those at the distance site. It is important to stress that conclusions about localization ability are dependent on the genetic structure of the population and the number of individuals examined. Increasing the number of individuals would be expected to increase the localization ability, if only due to obtaining replicate phenotypic measures on animals with identical local genotypes. Although an average localization ability of on the order of a megabase could be viewed as a disappointment, this is likely what is being achieved in other diversity outbred populations. Furthermore, if 1Mb is ∼1cM, localization ability is at least an order of magnitude better than what could be achieved in a single generation mapping experiment using a comparable number of animals (Lander and Botstein 1989).

**Figure 4:**
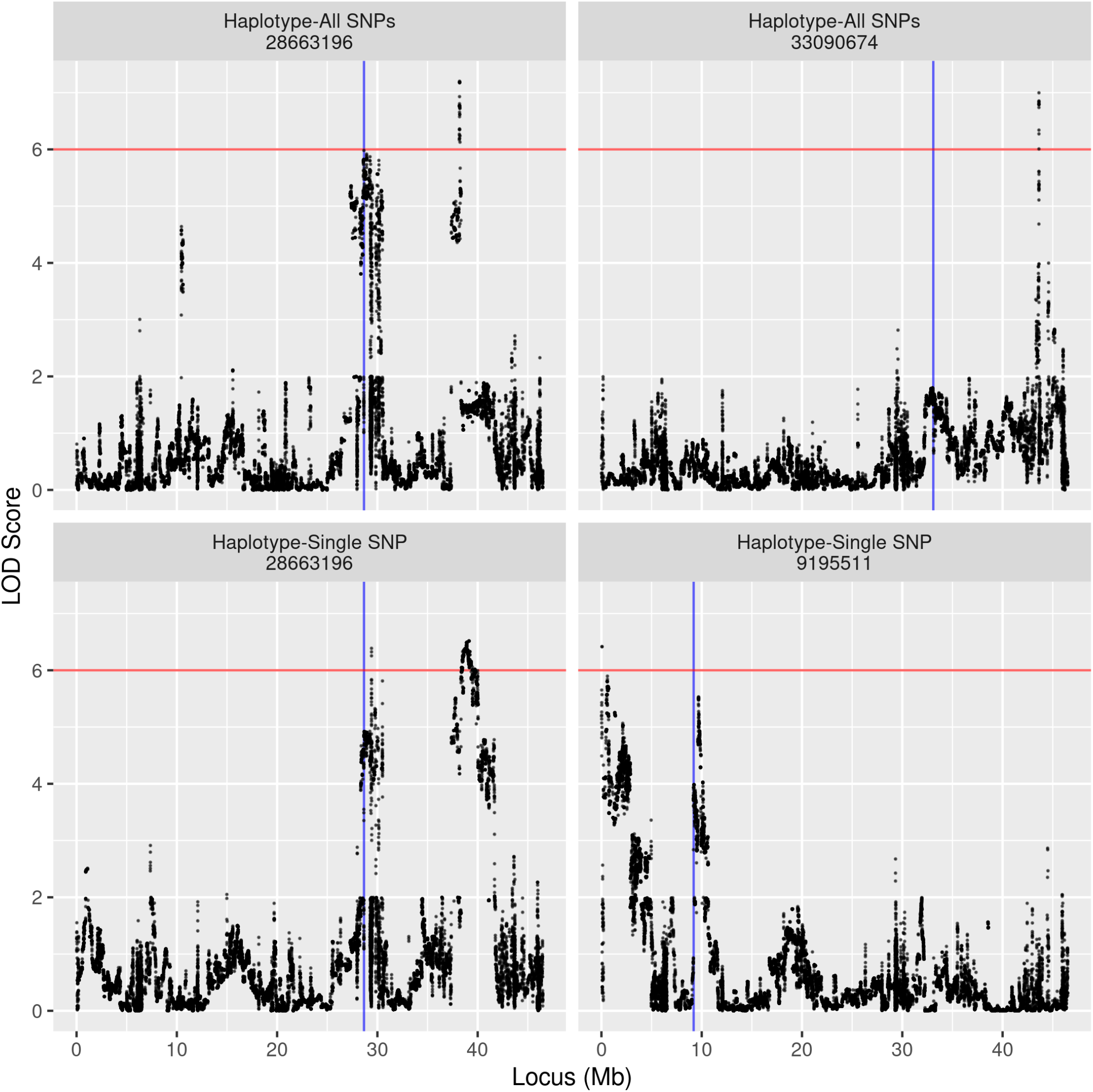
Four scans in which the most significant marker is located at great distance (> 7.5 Mb) from the causative gene. A red line indicates the significance threshold; a blue line is the location of the causative gene.

### A Demonstration Trait

A subset of 297 of the genotyped animals were assayed for bleeding time as a demonstration trait. This subset of animals for which a phenotype was measured drove the above decision to examine power in this subset of animals (the larger set of animals also included some close relatives). Bleeding time itself is not of great interest and does not appear to be highly heritable, but was easily measured on each animal at the same time a tail clip was carried out to obtain DNA for genotyping. Bleeding time is the number of minutes until blood clotting occurs following a tail clip. The vast majority of animals successfully clot in a few minutes with some taking 10 minutes or more. We transformed raw bleeding times to be normally distributed as we observed the trait to be highly skewd towards long bleeding times, and further removed the effects of sex, age (at time of assay), and date of birth from the trait before QTL mapping. We carried out both a marker- and haplotype-based genome scan on the residual normalized bleeding time as well as a shuffled set of the same phenotypes and employed thresholds for statistical significance derived above.

Supplementary Figure 5 are QQ-plots of genomewide LOD scores with actual bleeding time phenotypes relative to permuted phenotypes. There is evidently very little signal in these data as the QQ-plots are close to the unity line. This suggests the individual genes contributing to the trait have effects of less than the 5% per gene we consider in the simulations for an experiment examining ∼300 colony animals. Nonetheless regions with higher LOD scores are stronger candidates for harboring causative genes. Supplementary Figure 6 is a Manhattan plot of GWAS on permuted bleeding time (a negative control). Although no tests exceed the threshold for significance for either test, several marker-based test regions exceed 5, and a handful of regions exceed 4 for the haplotype-based test. Consistent with simulations and earlier claims when marker- or haplotype-based tests show “peaks” they do so via large blocks likely greater than 1Mb is size. Figure 5, a Manhattan plot of the bleeding time GWAS, shows no significant hits (consistent with the QQ plot). Furthermore it is not obvious from a visual comparison of the actual GWAS or GWAS on permuted phenotypes that the real GWAS is producing highly meaningful results. So any interpretation of the results must be tempered by the idea that observed peaks are only suggestive.

**Figure 5:**
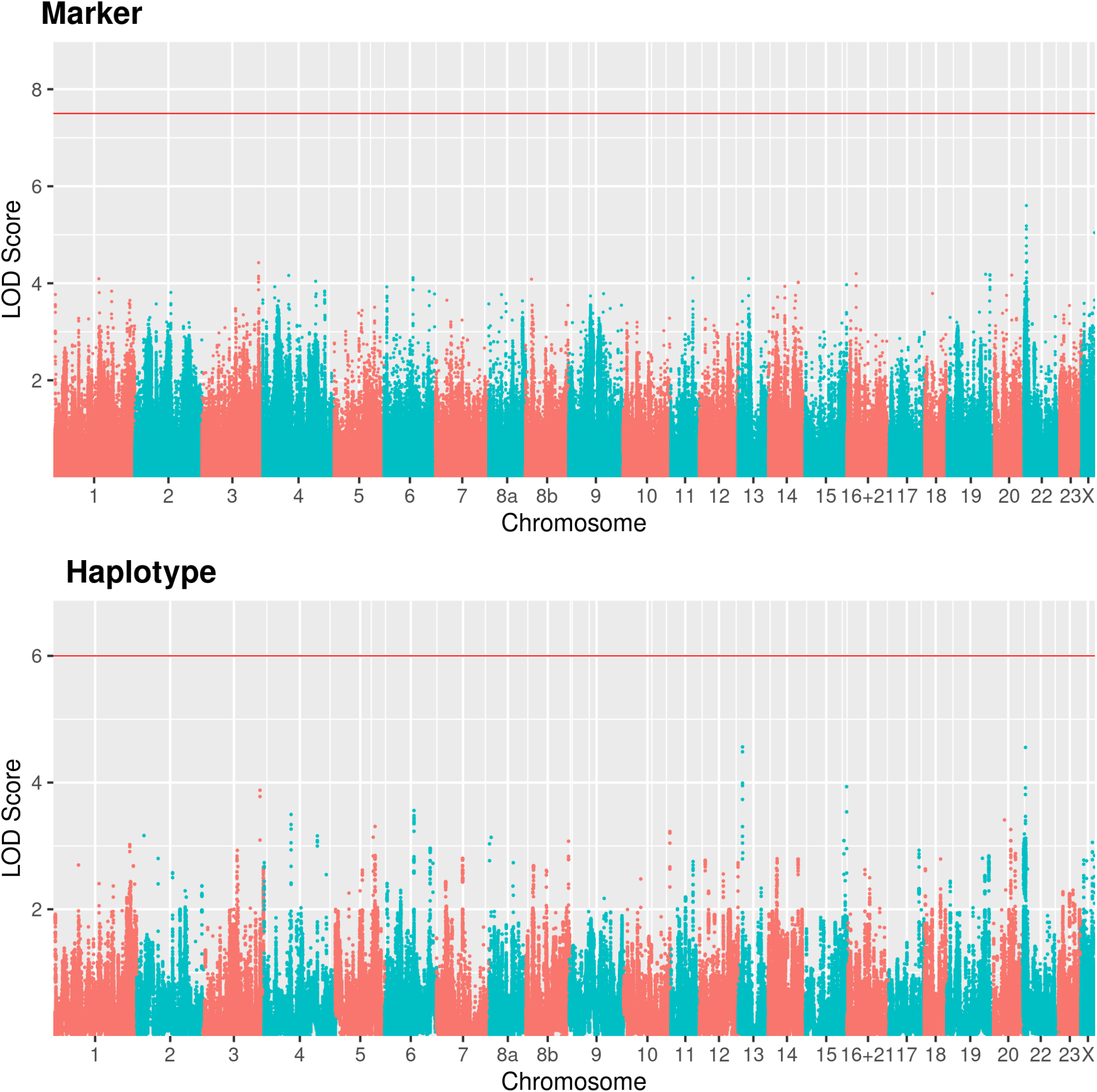
Manhattan plot of genome-wide scan for the bleeding time trait. Upper, marker-based scan; lower, haplotype-based scan. A red line indicates the significance threshold.

We choose to focus on four regions located on chromosomes 3, 13, 15, and 22 (Table 2). Although not significant, these peaks are elevated, show consistent signals across the two tests, show the expected block-like pattern of significance, and are not located near the tips of chromosomes (nor on the X chromosome) where imputation is more suspect. Supplementary Figure 7 shows the associations under the two tests for these four chromosomes and Figure 6 10Mb regions centered on the peak. Although the peaks are not significant they are consistent across tests and suggest localizations within about 2Mb. We thus tested a set of 83 genes whose mouse GO term matched “platelet function” and two additional genes associated with warfarin resistance in rodents to see if they were located within 2Mb of the candidate peaks. The peak on chromosome 3 was associated with three coagulation candidate genes, namely: Von Willebrand factor (VWF), CD9 antigen, and protein tyrosine phosphatase, non-receptor type 6. There are no obvious polymorphisms in VWF that could explain our mapping result, but it is an interesting candidate gene. We further hypothesized that variation in VKOR1 (or a paralog) could be associated with bleeding time QTL, as deermice are undoubtedly exposed to warfarin “rat” poison, but neither gene was associated with the four suggestive peaks.

**Figure 6:**
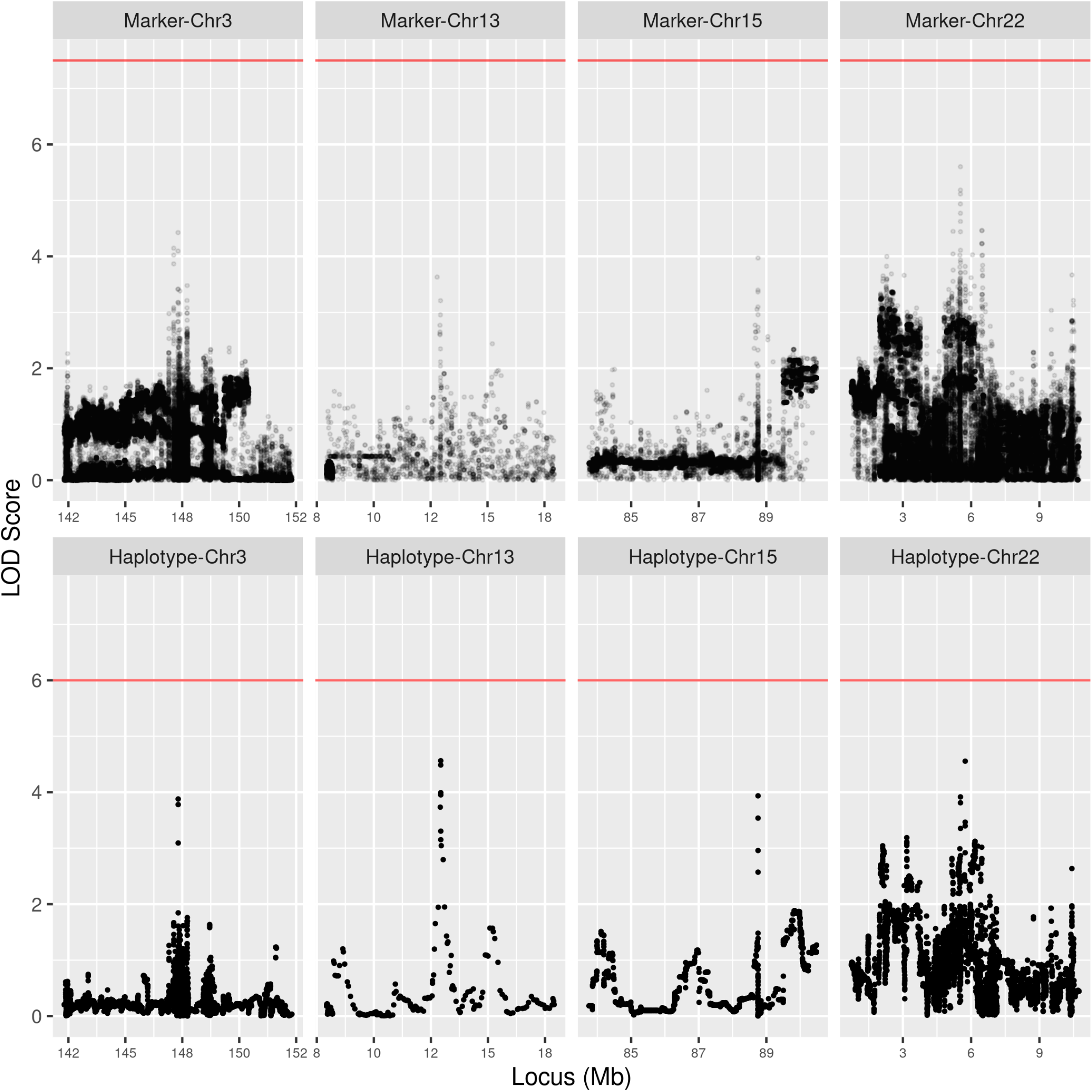
10 Mb regions centered around notable peaks from the genome-wide scan for bleeding time gene. Marker-based scans above and haplotype-based below depict the same regions on chromosomes 3, 13, 15, and 22. A red line indicates the significance threshold.

**Table 2:** Suggestive bleeding time hits

| Chr. | MSM |  | LOD |  | Candidate Gene (symbol:midpoint) |
| --- | --- | --- | --- | --- | --- |
|  | Marker | Haplotype | Marker | Haplotype |  |
| 3 | 147318730 | 147319452 | 4.03 | 3.21 | VWF:145711094, CD9:145856162, PTPN6:146604334 |
| 13 | 12785870* | 12931925 | 4.21 | 4.57 |  |
| 15 | 88753938 | 88753903 | 3.44 | 3.13 |  |
| 22 | 6471917 | 5738952 | 4.66 | 3.80 |  |

\* local MSM in 10 Mb region centered around haplotype-based MSM

## DISCUSSION

Here we explore a diversity outbred approach for dissecting complex traits in *Peromyscus leucopus*, a non-traditional genetic model system. We take advantage of a colony of deer-mice founded from 38 individuals between 1982-85 and subsequently maintained as a closed colony for 35+ years (∼40-60 generations). We speculate that this *P. leucopus* colony shares many genetic features with DO/HS mice and rat populations. Most importantly, by virtue of its large number of founders the colony segregates many alleles per gene, with the many generations of colony maintenance having broken down the initial linkage disequilibrium associated with the colony founding while also purging some of the initial diversity. Unlike DO/HS mice and rats, where the strains used to found the panel are both known and highly characterized, the *P. leucopus* colony founders are both unknown and tissues were never archived. As a result the DO mapping strategy of inferring diploid founder of origin haplotypes from marker data, and regressing on estimated haplotype dosages cannot be employed. Instead phenotypes must be regressed onto a set of genomewide imputed SNPs or local haplotypes inferred directly from the sample of genotyped individuals. Despite these potential shortcomings, we show that founder-unknown closed colonies harbor considerable mapping information that can be exploited to dissect complex traits.

We carried out low-pass sequencing on 405 mice (averaging ∼1X per animal) and used Stitch (Davies *et al*. 2016) to impute diploid genotypes at 17,751,882 SNPs. We took advantage of six deer-mice also characterized via RNAseq to validate Stitch-based imputations and show that imputed genotypes are generally very accurate. For the majority of mice we estimate 95% of the SNPs have errors in their gene dosage estimates of less than 0.03. We then carried out simulations utilizing a subset of 297 genotyped mice where we simulated QTL under three different genetic models and carried out chromosome 23 association scans using both a marker- and haplotype-based statistical test. This simulation approach is based on the idea of taking a set of individuals (or RILs) with high quality genotypes, simulating a complex trait conditional on the observed genotypes of these individuals, and then examining the ability to both detect and localize a simulated QTL (c.f.(McMullen *et al*. 2009; Aylor *et al*. 2011; King *et al*. 2012; Gatti *et al*. 2014)). This simulation strategy has the attractive advantage of grounding the patterns of genetic variation, linkage disequilibirum, and relatedness between individuals in reality. Conditional on the 297 deermice examined, the power to detect a QTL contributing 5% to a complex trait was ∼15-25% using a threshold that holds the false positive rate at one false positive per six genome scans. For the subset of simulation realizations with a significant hit we were able to localize the causative site to ∼2Mb-sized windows. Although power and localization ability differed among tests and genetic models, it did so only subtly.

A clear short-coming of the simulation approach employed, is that it is not simple to simulate additional individuals from the colony in order to examine power and localization in a larger experiment. It is perhaps obvious that increasing the number of deermice examined will increase both power and localization ability, but quantitative estimates are not available. If the output of Stitch were phased diploid genotypes, as opposed to unphased dosage estimates at each SNP, then it would be possible to simulate recombined chromosome-wide haplotypes to create additional pseudo-individuals, although such an extension is not trivial. A pure Monte Carlo approach to simulating colony animals from first principles could be attempted, but simulations might not fully encapsulate variation in (unknown) founder haplotype frequencies nor longer range within chromosome patterns of LD. Despite the power and localization ability of our approach being modest, it appears to be comparable to those seen in diversity outbred mice and rats *conditional on sample size*. In fact, a study of power in DO mice populations shows almost identical power to the estimates of this paper (c.f Figure 4 of (Gatti *et al*. 2014)), suggesting that like DO mice on the order of 500-1000 mice are likely necessary to routinely map QTL contributing 5% to variation in the complex trait (Chitre *et al*. 2020). Finally, QQ-plots suggest a great deal of signal relative to control scans, and a visual examination of Manhattan plots shows consistent signal associated with simulated QTL locations which would be expected to grow with larger sample sizes. Thus despite the founders of the *P. leucopus* colony being unknown, colony animals can be utilized much the same way as DO mice or rats to dissect complex traits in *P. leucopus*.

We explored three genetic models and two statistical tests. Although one of our models simulates the customary single SNP contributing ∼5% to variation in a complex trait, in light of human GWAS studies generally failing to identify intermediate frequency alleles of large effect (Manolio *et al*. 2009), this model is almost certainly wrong in most cases. We thus chose to simulate two additional models in which variation at a gene contributing to a complex trait was due to ten or potentially hundreds of causative SNPs in that gene. Such models are broadly consistent with the idea of mutation selection balance maintaining variation at complex trait genes (Pritchard 2001; Thornton *et al*. 2013). Although these simulations are undoubtedly an over-simplification of such multiple causative site models, they may be useful for comparing marker-versus haplotype-based tests. We hypothesized that single marker tests would perform well under the single causative site model (but not necessarily multi-site models), whereas haplotype-based tests would perform well under multiple causative site models. But, we observe that both marker- and haplotype-based statistical tests performed similarly (based on QQ plots) irrespective of the underlying genetic model. The megabase sized blocks of elevated marker-based LOD scores associated with the location of simulated QTL suggests that LD is extensive enough in colony mice that several hundred SNP-markers are capable of tagging causative regions.

We finally carried out a GWAS on a demonstration trait, bleeding time following a small tail clip, and did not observe any regions exceeding our genome-wide false positive controlling threshold. Nonetheless we looked more carefully at the four top scoring regions showing consistent signal over tests and in two cases the peaks are within 2Mb of platelet function candidate genes. Although this degree of co-localization is not inconsistent with pure chance, it is of interest that one of the candidate genes is Von Willebrand factor (VWF), a gene at which mutations in humans result in a well-studied blood clotting disorder. We have not carried out functional validation of VWF, but a VWF knock-out in the mouse exhibits a prolonged bleeding time (Ni *et al*. 2000), highlighting a potential weakness of current knockdown-based functional validation experiments involving co-located candidate genes.

Although *Peromyscus* is not considered a genetic model system, the genus is well suited for addressing several questions of biologist interest including the genetic bases of longevity, behavior, physiology, adaptation, and a deermouse’s ability to serve as a disease vector. Each of these phenotypes are associated with clear scientific and/or human disease questions where the ability to dissect complex traits in *Peromyscus* would be of great value. Our results suggest that individuals from a long-maintained colony of *P. leucopus* deermice can be utilized in a manner very similar to DO/HS mice and rats for dissecting complex traits. We simulate 300 individuals in this study, motivated by the number of individuals for which we measure a demonstration trait, but work here shows that much greater than 300 mice must be phenotyped and genotyped to consistently dissect complex traits.

The results of this study are not only of interest to *Peromyscus* researchers. The simulations of this work show that virtually any closed colony of animals derived from a limited number of founders and maintained for several dozen generations can be used to dissect complex traits using a DO/HS-style approach. This can be accomplished cost effectively using low pass short read sequencing and genotype imputation provided the species exhibits high levels of nucleotide diversity and has a reference genome. Colonies derived from unknown founders clearly have some disadvantages, but an unknown-founders colony is far from a complete impediment. Given an annotated genome and a GWAS scan, candidate genes can be identified, and the genetic structure of the colony characterized with respect to founder haplotypes for only small regions of interest. It is also possible sophisticated extensions to Stitch (Davies *et al*. 2016) will eventually allow for phased haplotype estimates from large samples from closed colonies. Our work should further motivate a founder unknown DO/HS-strategy for a large number of species having extant closed colonies.

## ACKNOWLEDGEMENTS

Work was funded by NIH grants R21AI126037 and R01AI57513 to ADL and AGB. This work was made possible, in part, through access to the Genomics High Throughput Facility Shared Resource of the Cancer Center Support Grant (P30CA-062203) at the University of California, Irvine and NIH shared instrumentation grants 1S10RR025496-01, 1S10OD010794-01, and 1S10OD021718-01.

## COMPETING INTERESTS

No competing interests declared.

Table 1: Properties of complex trait mapping simulations.

---

**Supplementary Figure 1:**
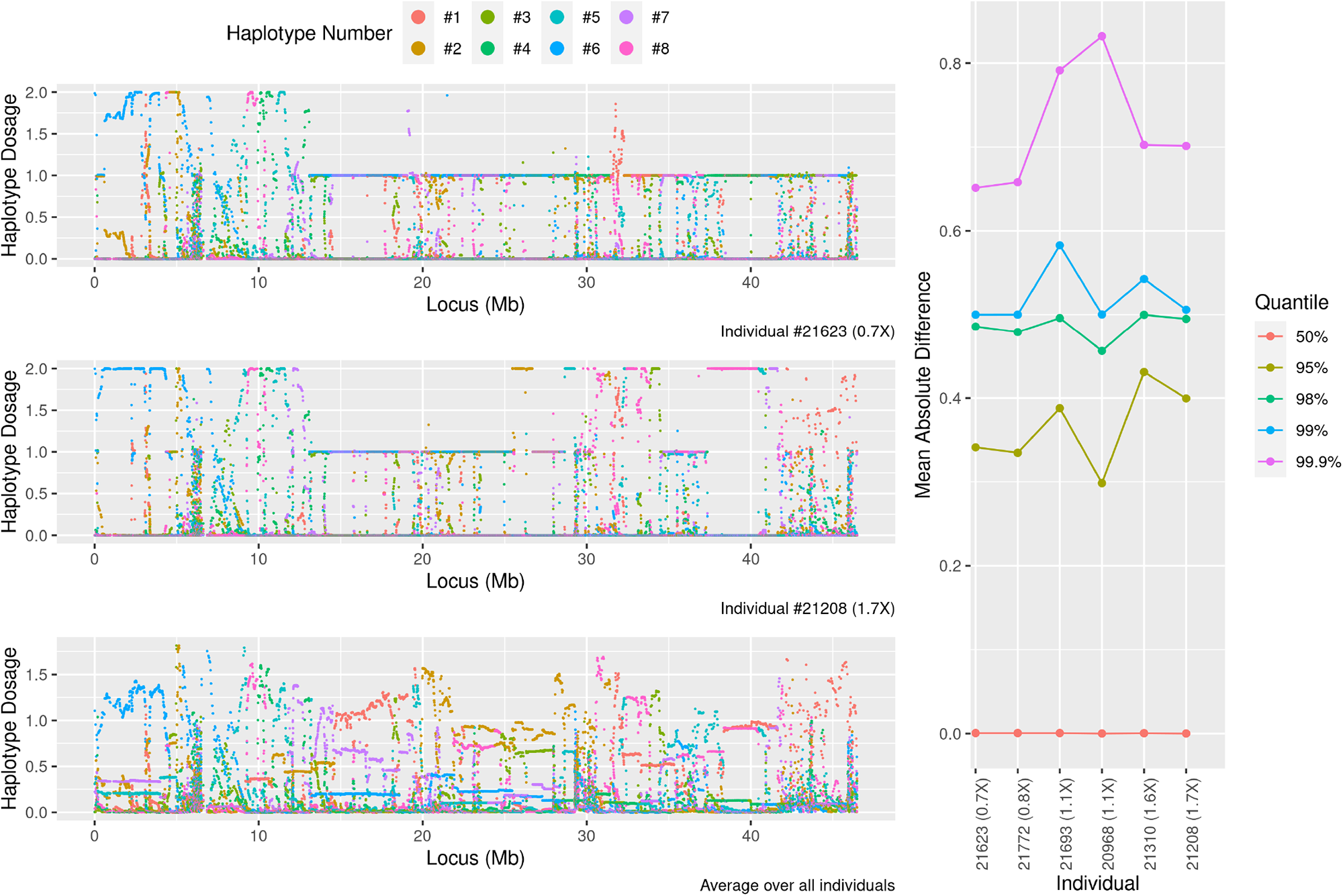
Stitch imputed haplotypes. Left panels from top to bottom are raw dosage values colored by haplotype for a lower coverage individual (0.7X), higher coverage individual (1.7X), or average over all 297 individuals. Right panel is the mean absolute change in every tenth haplotype dosage scores for 6 individuals.

**Supplementary Figure 2:**
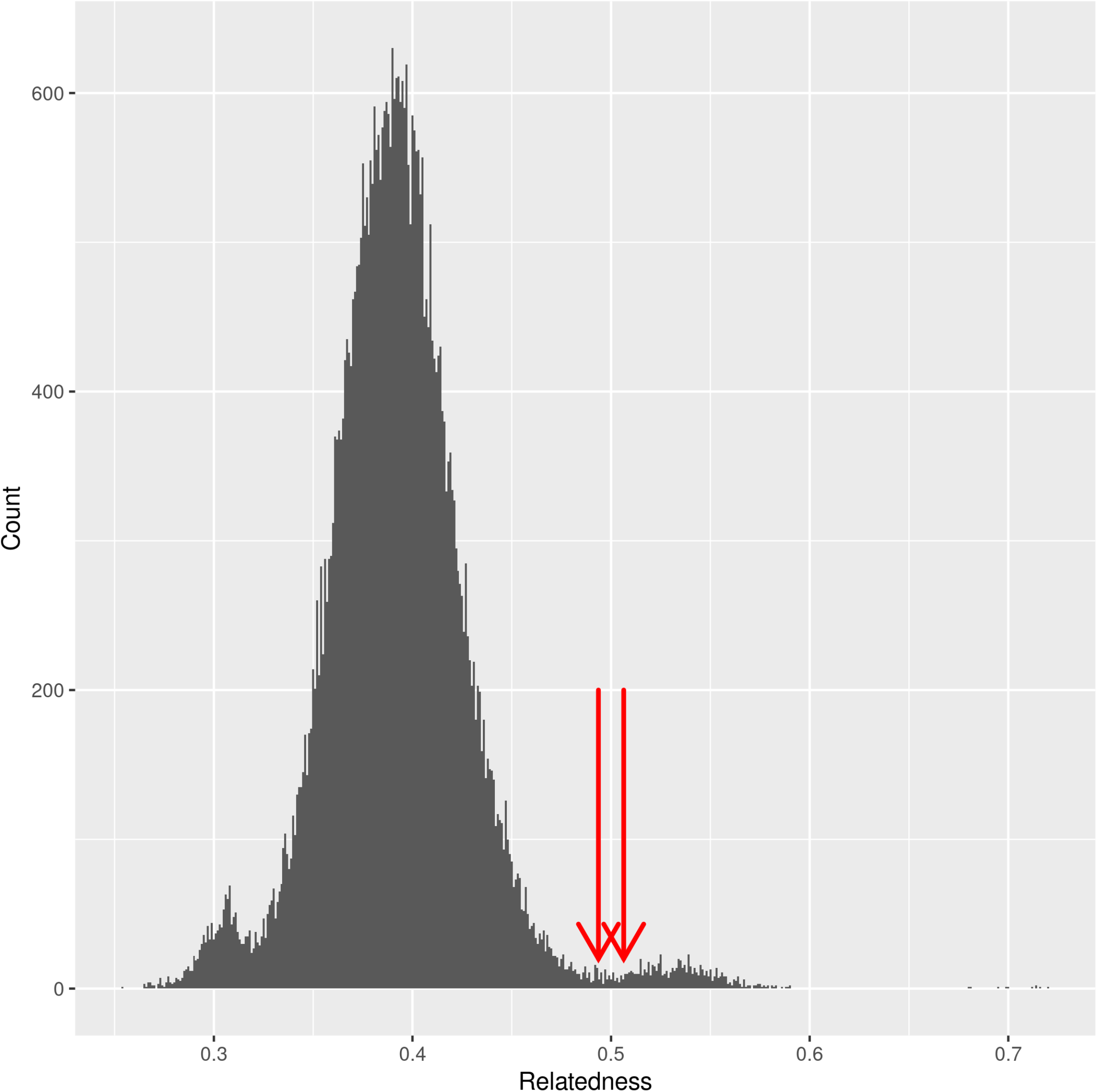
Histogram of relatedness values in the kinship matrix used in QTL scans. Two arrows point to relatedness values of known parent-offspring relationships.

**Supplementary Figure 3:**
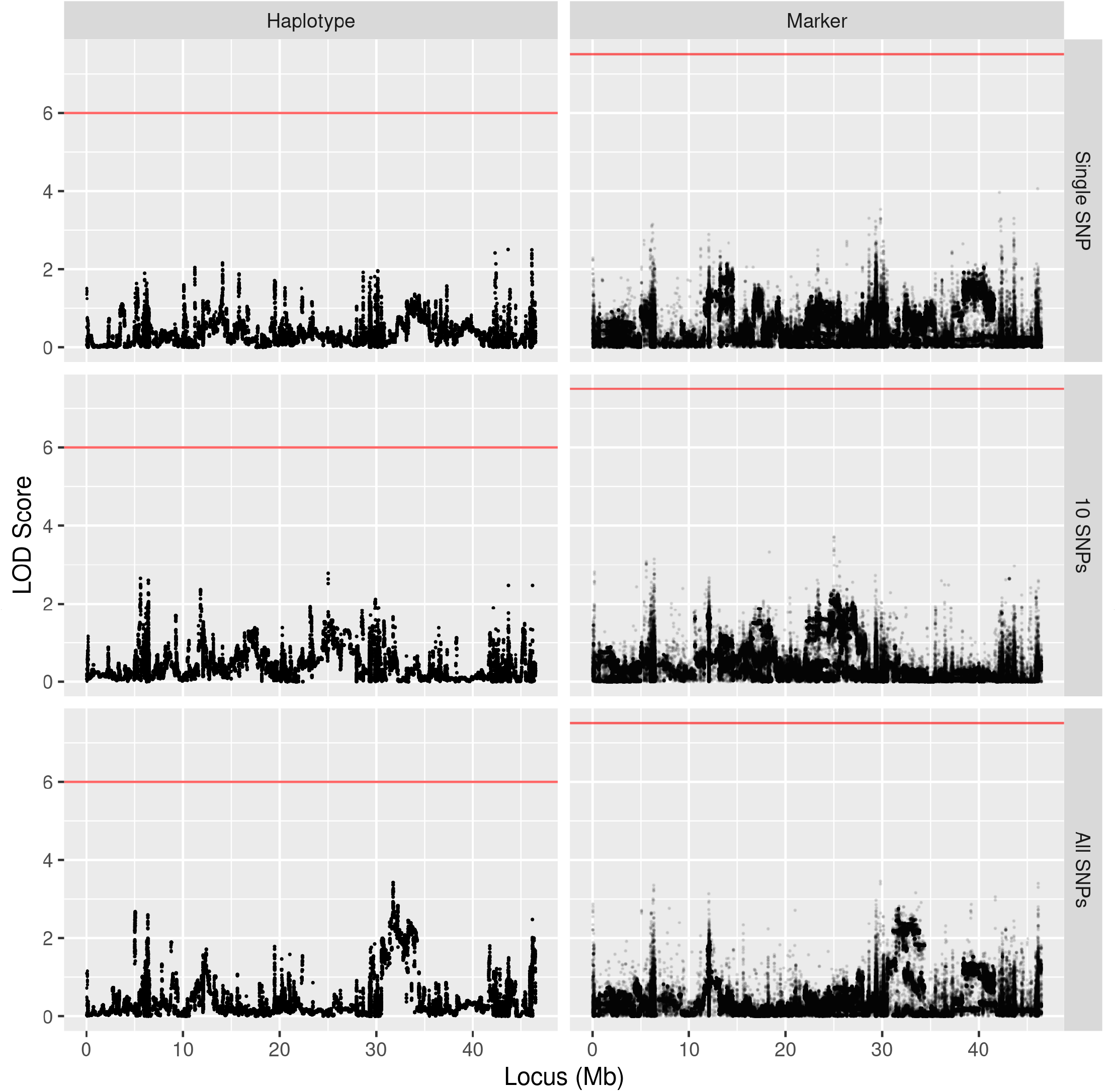
Example of chromosome 23 scans with a simulated QTL on chromosome 19 (control). Faceted to match Figure 3.

**Supplementary Figure 4:**
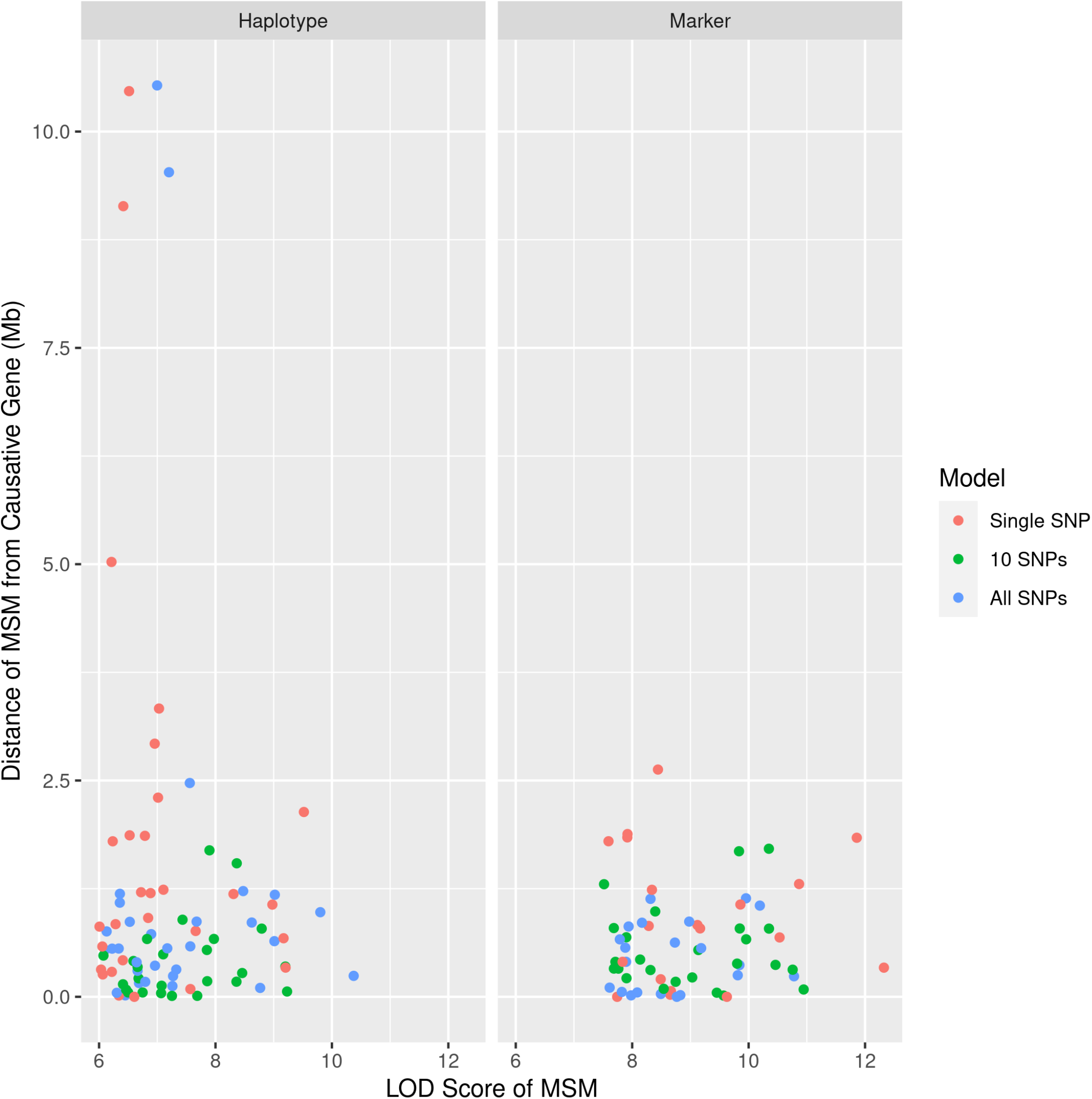
Distance of most significant marker (MSM) in a scan from the simulated causative gene as a function of the MSM LOD Score. Only replicates having one or more “hits” are displayed. Left, haplotype-based, right, marker-based.

**Supplementary Figure 5:**
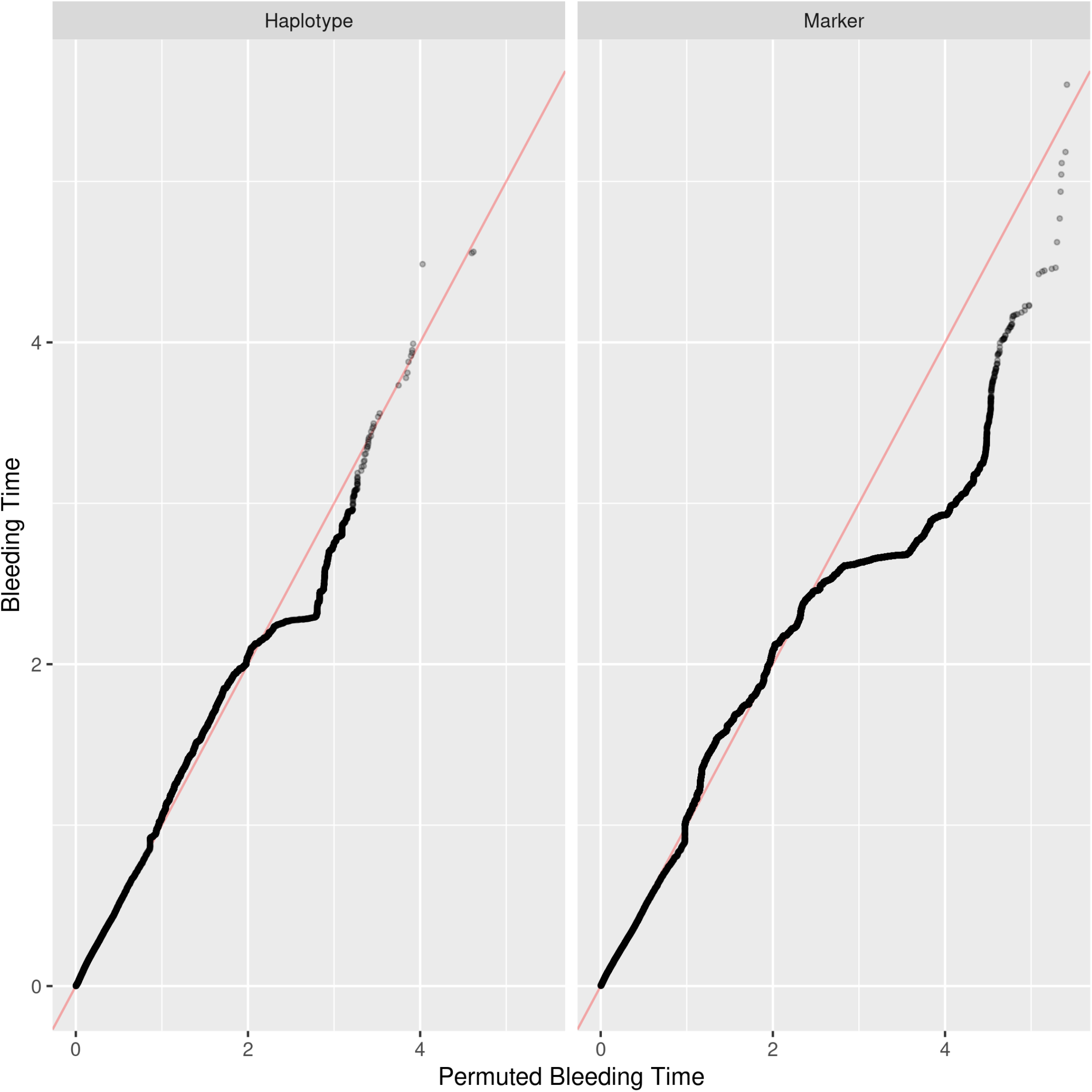
QQ plots of LOD Scores from genome-wide scans for bleeding time with actual or permuted phenotypes. Left, marker-based tests; right, haplotype-based tests.

**Supplementary Figure 6:**
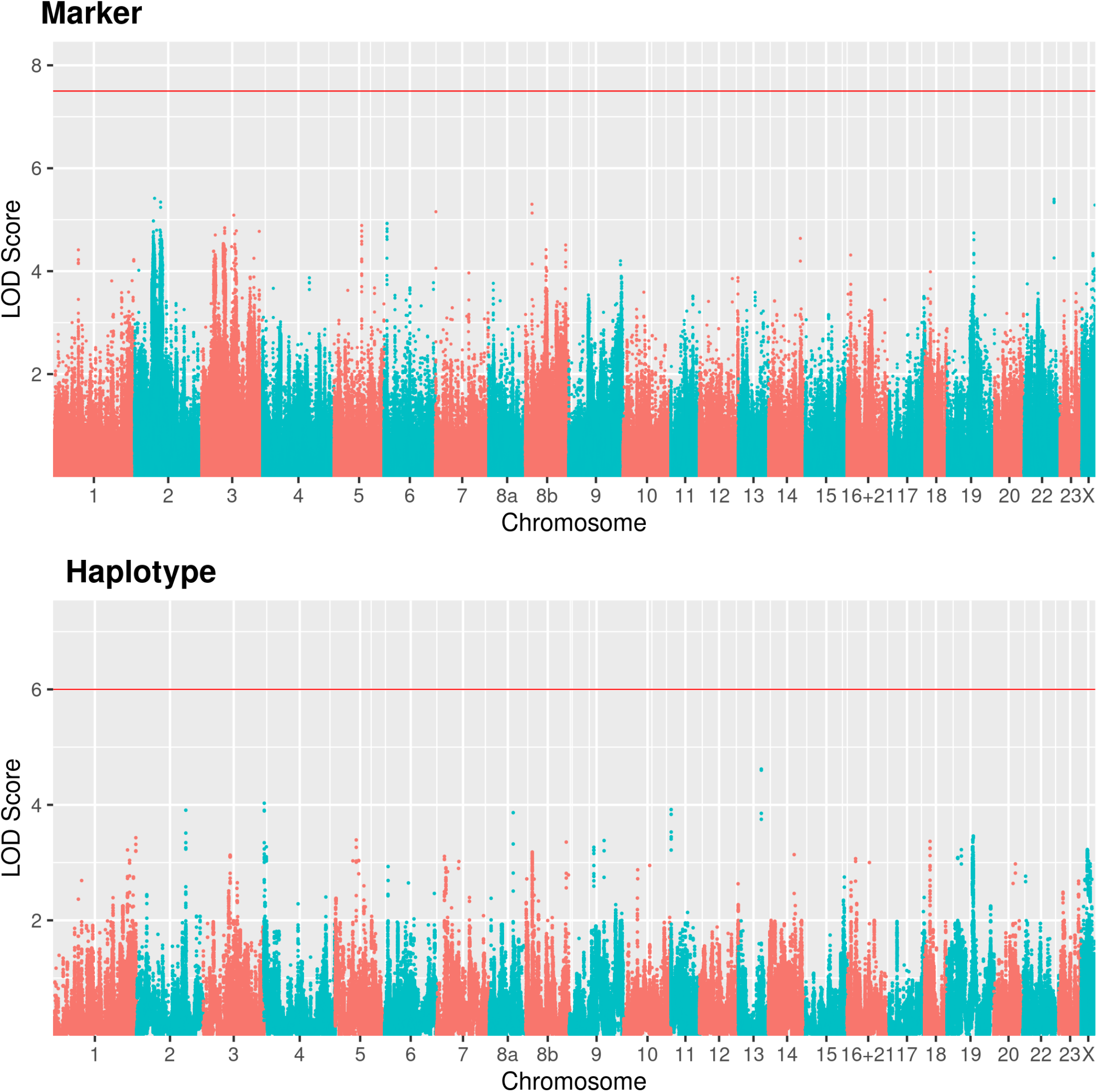
Manhattan plot of a control genome-wide scan for bleeding time under permuted phenotypes (negative control). Faceted to match figure 5.

**Supplementary Figure 7:**
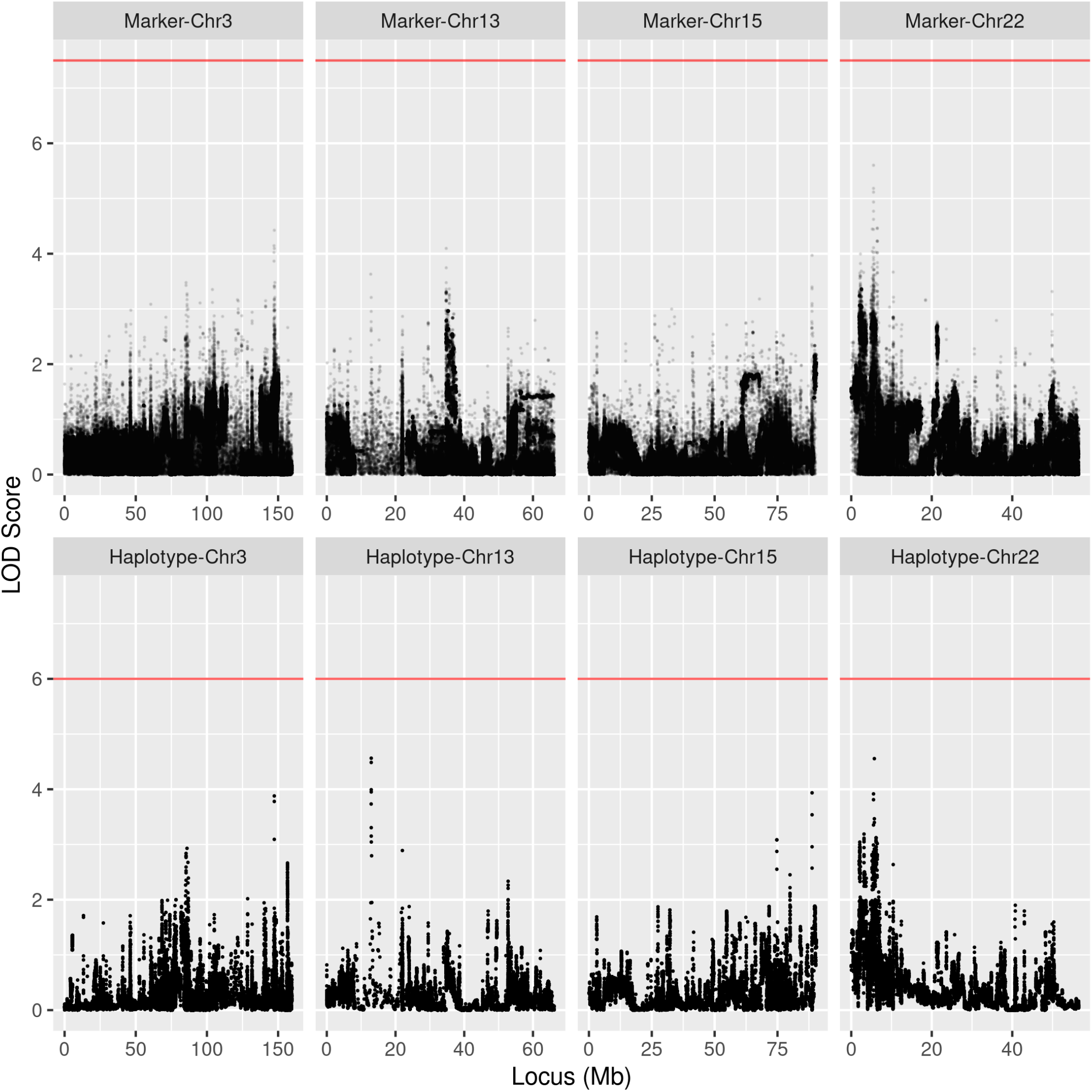
Chromosome scans of chromosomes containing notable peaks from the genome-wide scan for bleeding time gene. Faceted to match figure 6.

